# UVC Inactivation of Black Mold is Wavelength-Dependent, and its Growth in HVAC Systems is Preventable Using Periodic Dosing with commercially available UVC LEDs

**DOI:** 10.1101/2022.06.06.495021

**Authors:** Richard M. Mariita, Rajul V. Randive, Michelle M. Lottridge, James H. Davis, Benjamin W. Bryson

## Abstract

Mold growth in HVAC systems poses a threat to human health, increases facility management operating costs due to decreased air flow efficiency, and damages buildings, impacting buildings’ attractiveness in an ever more competitive property market concerned with both carbon footprint and ventilation quality. Although UVC treatment of HVAC systems has been shown to reduce growth, efficient design requires a specified amount of UVC dose coupled with a specific dosing strategy. UVC disinfection efficacy and wavelength sensitivity of spores from the most common and notorious black mold, *Cladosporium halotolerans* is not known. This study investigates the sensitivity of *C. halotolerans* and demonstrates that growth of black mold on HVAC coils can be effectively prevented with a periodic dosing scheme using commercially available UVC LEDs. Multiple UVC LED arrays with varying wavelength peaks in the range of 252-280 nm were used to demonstrate the spectral sensitivity of *C. halotolerans* by keeping the wavelength specific UVC dose equal in order to avoid bias. The data obtained for doses of 25, 75, 125, 175, and 225 mJ/cm^2^ was used. The data obtained was used to determine a dose response curve and susceptibility patterns. Disinfection performances for the arrays were determined using log reduction value (LRV) by comparing the controls against the irradiated treatments. To study mold growth prevention on HVAC coils, the 267 nm array which had the best disinfection efficacy was utilized. The coils were placed in a chamber with a temperature set at 24 °C and relative humidity (RH) at 96%. Unlike the controls, the test coil was irradiated with a dose of 28.8 mJ/cm^2^ every 12 hours. The coils were monitored, and observations were recorded using time-lapse videos. The highest disinfection level of black mold observed in the range of 250-280nm occurred at 267 nm, with 225 mJ/cm^2^ obtaining 4.03 LRV. Linear regression analysis at 95% for 252 nm (R^2^ =0.9637, *p*=0.0030), 261 nm (R^2^ =0.9711, *p*=0.0021), 267 nm (R^2^ =0.9723, *p*=0.0020), 270 nm (R^2^ =0.9819, *p*=0.0010), 273 nm (R^2^ =0.9878, *p*=0.0006), and 280 nm (R^2^ =0.9914, *p*=0.0003) displayed significant association between arrays’ peak wavelengths and disinfection performances against *C. halotolerans*. The study revealed that a higher dose of 225 mJ/cm^2^ is required to disinfect *C. halotolerans* by 99.99% (4.03 LRV). However, using a periodic dosing strategy utilizing 28.8 mJ/cm^2^ prevented any mold growth, while the fungal levels in the positive control increased. The study allows for the design and implementation of mold growth prevention strategies in HVAC systems to improve health and lower operating costs.

## Background

Mold and poor indoor air quality are a public health concern because it affects the physical, mental, and social health of the residents in shared enclosed spaces [1] where they spend 90% of their time [2]. Mold can for example cause allergic reactions in unprotected workers, first-responders, homeowners, and volunteers in recovery and restoration of moldy indoor environments after hurricanes, typhoons, tropical storms, and flooding damage which are a growing concern for healthcare providers and disaster medical practitioners throughout the world [3]. Some toxic molds produce mycotoxins, which may cause disease when inhaled [4].

Historically, mercury-based UVC lamps have long been used for surface treatment within Heating, Ventilation, and Air Conditioning (HVAC) systems [5]. Target UVC disinfection areas include coils, drain pans, plenum walls, humidifiers, fans, and filters [6] in systems installed in buildings and automobiles [7]. Mold growth prevention is important as it can colonize heating, ventilation, and air conditioning (HVAC) systems leading not only to restriction of airflow [8], but its presence can lead to the production of volatile organic compounds (VOCs), responsible for the poor quality of air [9]. Additionally, inactivated/dead mold can still cause allergic reactions [10]. Thus mold growth prevention in HVAC systems is a better strategy than disinfection.

Common strains of interest throughout nearly all regions of the world are black molds of the *Cladosporium* Genus. Moreover, *Cladosporium halotolerans* is the most frequently isolated *Cladosporium* species indoors [11], [12] and can be found in marine, freshwater, and terrestrial environments [13] and even on human scalps [14]. In particular, *C. halotolerans* has been isolated in the US, China, Europe, New Zealand, India, Taiwan, Africa, and South America [11]. It also survives humid and salty conditions much better than other strains and thrives in areas such as bathrooms [13]. A survey of select locations in air handling units (AHUs) has shown that *Cladosporium* sp. fungi are not only the commonly recovered in terms of growth sites but also deposits spores in the blower wheel fan blades, the ductwork, and the cooling coil fin [15]. One study has revealed that even within the *Cladosporium* Genus, *Cladosporium sphaerospermum* species complex (where *C. halotolerans* belongs to) are most predominant representing 44.7% [16]. The *C. sphaerospermum* species complex was found to contribute 23.1% of indoor air and 58.2% of indoor surfaces isolates [16]. With lower water activity (≥ 0.82) (Figure 1) compared with the other *Cladosporium* complexes, the xerotolerance of *C. sphaerospermum* species complex offers an advantage of colonizing ambient indoor environments [16]. Although not as widespread as *Cladosporium* sp., other molds that have been recovered from cooling coil fins and insulation include *Penicillium* sp., *Aspergillus* sp., and *Paecilomyces* sp. [15].

**Figure 1.**
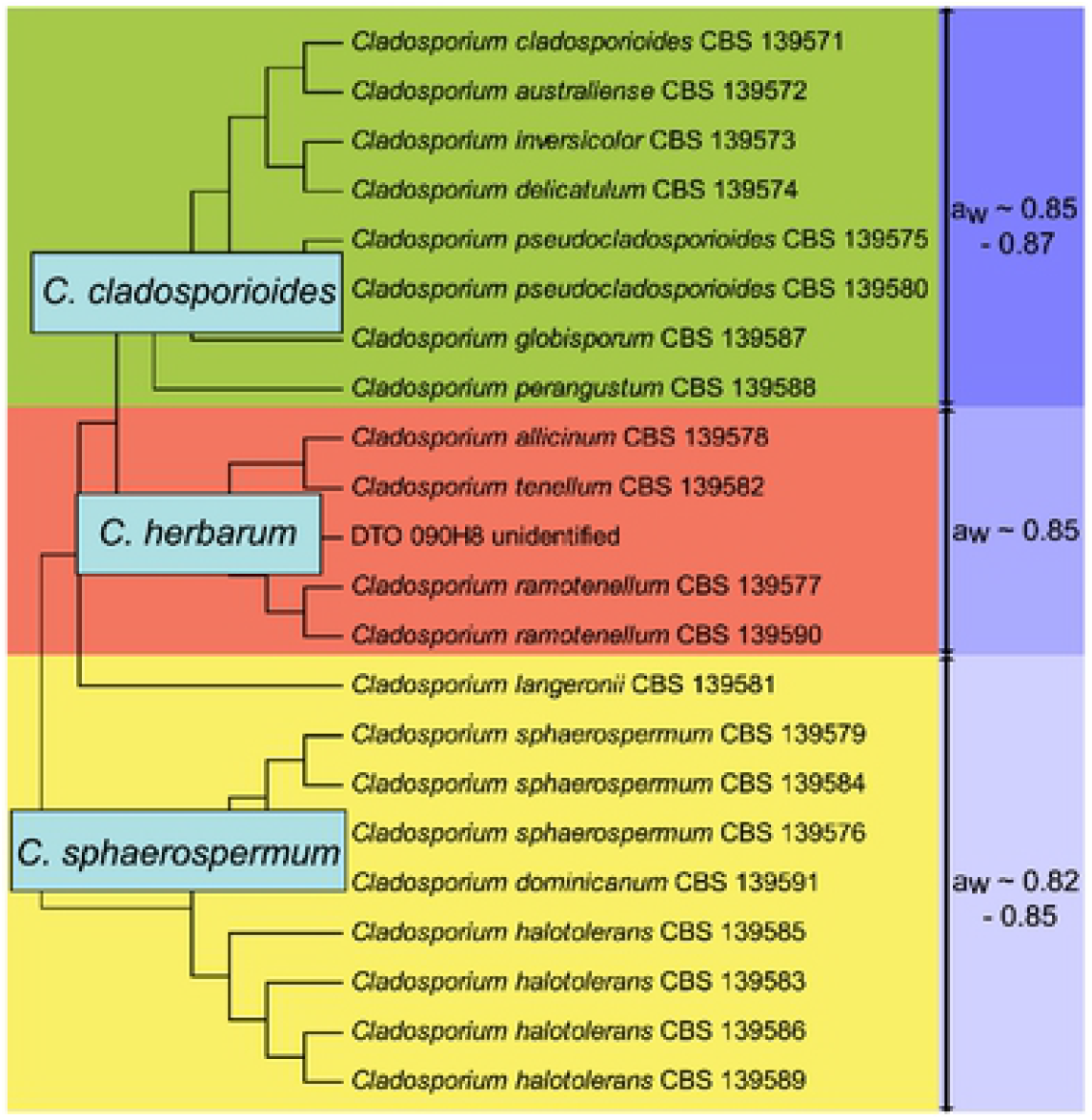
Schematic dendrogram 22 Cladosporium species and their minimum water activities supporting xerotolerance of *C. halotolerans* which enables it colonization of indoor spaces. Organisms arc grouped into three species complexes, *C. cladosporioides* (Green), *C. herbarum* (red), and *C. sphaerospermum* (yellow). Figure reused from Segers et al. [16] under Creative Commons (CC BY 4.0) license http://creativecommons.org/publicdomain/zero/l.0/ (Accessed on 25^th^ May 2022).

Sometimes frequently replaced filters, dehumidification, and switching to alternate air sources can be used to address mold growth and presence. However, the HVACs cooling systems invariably cause precipitation to form, while external air use can be inefficient and is not always viable, and filters can restrict air flow, fail to prevent all colonization, and do become colonized themselves. UVC lamps have been used to decrease and prevent mold growth on the moist interior surfaces of HVAC systems [17]. However, commercially available UVC LEDs offer a promising application replacement of mercury lamps, reduce energy consumption and replacement cycles by having a smarter and targeted periodic disinfection cycle, leveraging the instant on/off and no hysteresis characteristics of UVC LEDs.

To design a system effectively, it would be typical to estimate the amount of irradiation needed to achieve a certain disinfection or mold growth prevention on a surface dosed with UVC to assess the number, distance, and placement of UVC LED lights. However, for successful implementation,UVC sensitivity of critical fungi like *C. halotolerans* and *C. sphaerospermum* should be known. And despite extensive studies on wavelength sensitivities against bacteria such as *E. coli* [18], little information on the sensitivity versus wavelength for mold species including *C. halotolerans*. In this study, we assess the wavelength sensitivity of these molds with LEDs to facilitate future designs.

In some applications, it is possible to increase system lifetime and reduce cost, both in term of electrical consumption, replacement cycle and associated costs such as scheduling, recording, disposal of old mercury lamps, and inability to use the space during maintenance. Further cost reductions can be achieved by shifting from constant periodic dosing with UVC to eliminate microbes as soon as they enter the system to periodic dosing with UVC to prevent microbes from establishing in a system. Especially because at 25°C, *C. halotolerans* can grow at a rate of almost 4 mm/day [16]. Thus, preventing mold growth in HVAC systems via periodic dosing may benefit from such ventilation system designs. Prevention of mold growth in HVAC systems can help support the creation of energy-efficient dwellings; while protecting the health of the people in enclosed spaces from respiratory problems like, asthma, allergic rhinitis, and other respiratory pathologies [19]. Additionally, use of UVC as a supplementary addition to the existing mechanical ventilation offers additional benefits in the form of eACH [20].

## Methods

### Mold propagation and long-term storage

The strain *Cladosporium halotolerans* ATCC 10391 (NRRL 1671) was obtained from the American Type Culture Collection (ATCC). ATCC Medium: 200 YM Medium (Agar Broth) (pH 6.2). A pure colony was obtained and used to make mold stock. Incubation was done at 20-25°C for 2 weeks days before harvest [21]. Mold suspensions were made in glycerol sterile 10% (v/v) for long-term storage in a -80°C freezer by dispensing 0.5 mL aliquots into labeled cryovials.

### Black mold strain and growth conditions

Pure colonies were obtained from stock and used to grow test cultures. During incubation, each petri dish was individually wrapped using 3M micropore tape (Catalog No.19-027761). Test mold harvesting was done using ice-cold sterile 10 mM *N*-(2-acetamido)-2-aminoethanesulfonic acid (CAS-No. 7365-82-4), 0.02% Tween 80 (CAS-No. 9005-65-6) (ACES; pH 6.8) [22]. Mold was then standardized and further incubated for up to 1 week to determine disinfection inoculum size.

Incubation was done at 24°C and 97% RH. Temperature and humidity were monitored using HOBO® MX Temp/RH Logger (MX1101). Genera identification and confirmation of black mold was performed using bright microscopy (Figure 2) by observing verrucose conidia as a distinguishing feature [23]. General confirmation of mold was performed using bright microscopy with staining performed using 1.0 mg/mL propidium iodide solution (Thermo Fisher, Cat # J66584). Here, mold was stained and allowed to settle for 60 seconds before applying a cover slip and imaging done using Zeiss Axiovert inverted microscope.

**Figure 2:**
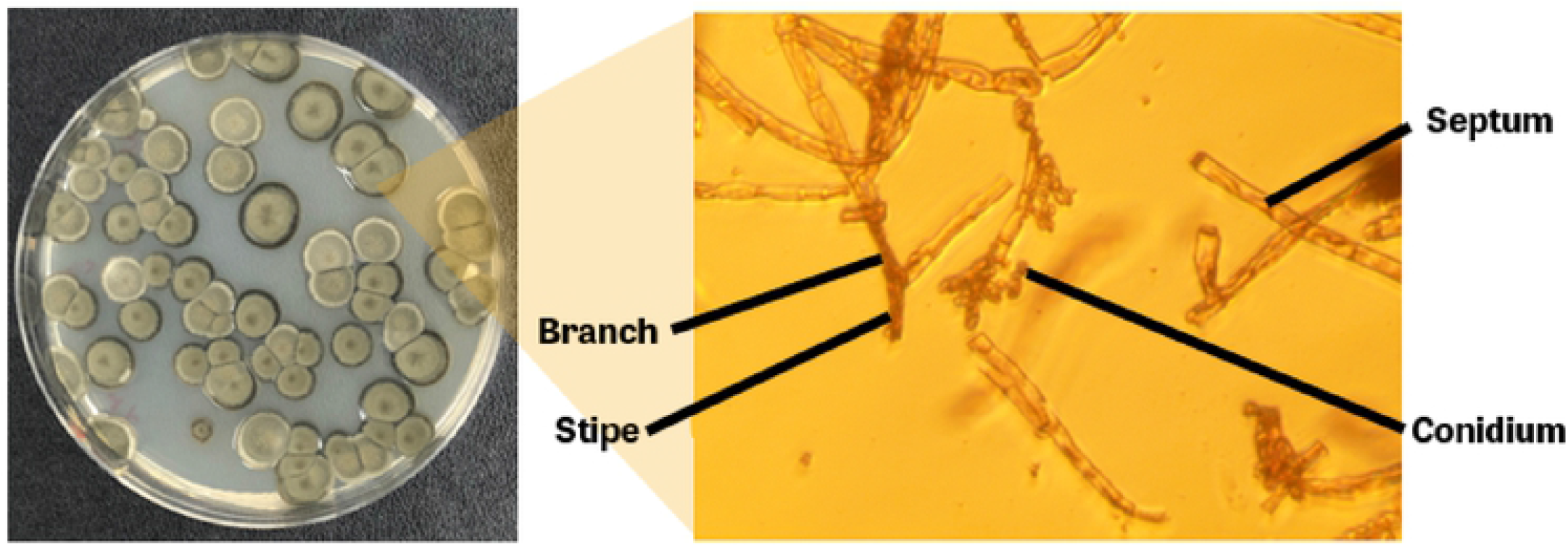
Visual inspection and observation of verrucose conidia (distinguishing feature) of *C. halotolerans* as observed under the light microscope was used for microscopic confirmation of the test organism. Bright-field microscopy was done on day 7 (stained using **Propidium Iodide**). Fungal hyphae for *C. halotolerans* ATCC 10391 are septated and observed at 100 X with immersion oil using a Zeiss Axiovert inverted microscope.

### Irradiance dose

Incident irradiance (fluence) was measured using an X1 MD-37-SC1-4 optometer calibrated to 265 nm. Because of the deviation of calibrated wavelength, it’s expected an error of 5% may be introduced from the optometer. The wavelength of the LEDs was confirmed using a Maya 2000 pro series spectrometer (Ocean Optics) (Figure 3). This study utilized a proprietary periodic dosing strategy [24] to prevent mold growth.

**Figure 3:**
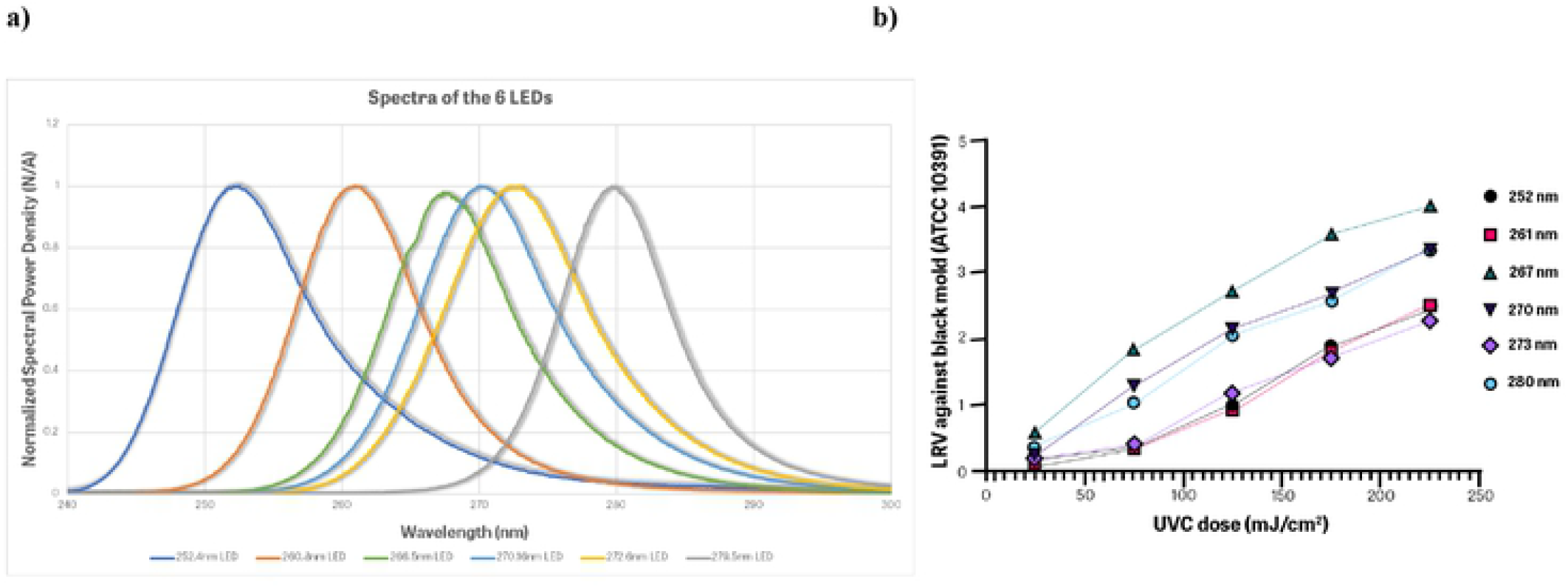
**a)** Peak wavelengths (252-280 nm) of the arrays used as confirmed using Maya pro 2000 (Ocean Optics) **b)** Wavelength sensitivity of black mold revealed most sensitivity at 267 nm

### Simulations

Simulations for this work were done using Zemax OpticStudio 22.1.1 (https://www.zemax.com/) using source data from a Klaran KL265-50V-SM-WD with a down-rated power output of 58mW to account for use case conditions. A total of 9 LEDs were used at a 1” (25.4mm) spacing to represent the board used in testing. The simulated board was centered above the representative coil and virtual detector at a 4” (103mm) height.

### Black mold growth prevention

The test and positive control coils (part number J1800002410) were nebulized with an equal amount of spores suspended in 10 mM *N*-(2-acetamido)-2-aminoethanesulfonic acid (CAS-No. 7365-82-4), 0.02% Tween 80 (CAS-No. 9005-65-6) (ACES; pH 6.8). The negative control coil was neither UVC dosed, nor treated with mold. The relative humidity of the test chamber was maintained at 97 % and the temperature was kept at 24°C.

### Statistical analysis

The determination of trends for each array as a function of performance was statistically analyzed using linear regression (simple exponential model). Additionally, an unpaired t-test (two-tailed) was used to measure the statistical significance of arrays’ disinfection performances. GraphPad Prism 9.1.2 (GraphPad Software, Inc.) was used to perform statistical analysis.

## Results

The study objectives were to determine the dose response for *C. halotolerans* ATCC 10391, as well as investigate the amount of dose required in a periodic dosing strategy to prevent mold growth on HVAC coils. Our results demonstrate that black mold is most sensitive to 267 nm, with an inactivation rate constant (susceptibility rate) of *k*= of 0.129388 (Table 1, and Figure 4). With a 267 nm peak wavelength array obtaining the fastest inactivation rate, better incremental log inactivation will be obtained compared to other tested peak wavelengths.

**Table 1:**
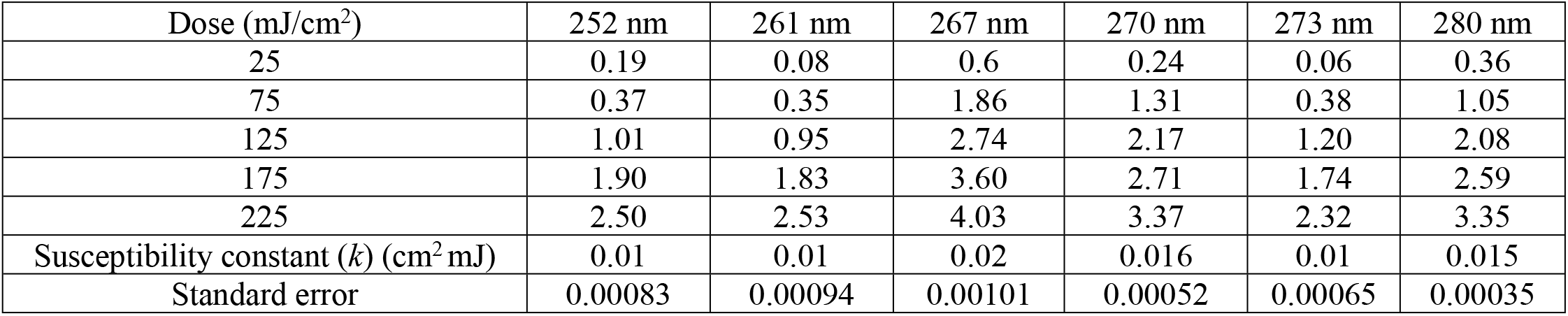
UVC efficacy against *C. halotolerans* with different wavelengths. The study revealed that with 267 nm peak wavelength yielded 4.03 LRV (99.99% black mold reduction) with 225 mJ/cm^2^.

| Dose (mJ/cm<sup>2</sup>) | 252 nm | 261 nm | 267 nm | 270 nm | 273 nm | 280 nm |
| --- | --- | --- | --- | --- | --- | --- |
| 25 | 0.19 | 0.08 | 0.6 | 0.24 | 0.06 | 0.36 |
| 75 | 0.37 | 0.35 | 1.86 | 1.31 | 0.38 | 1.05 |
| 125 | 1.01 | 0.95 | 2.74 | 2.17 | 1.20 | 2.08 |
| 175 | 1.90 | 1.83 | 3.60 | 2.71 | 1.74 | 2.59 |
| 225 | 2.50 | 2.53 | 4.03 | 3.37 | 2.32 | 3.35 |
| Susceptibility constant (*k*) (cm<sup>2</sup> mJ) | 0.01 | 0.01 | 0.02 | 0.016 | 0.01 | 0.015 |
| Standard error | 0.00083 | 0.00094 | 0.00101 | 0.00052 | 0.00065 | 0.00035 |

**Figure 4:**
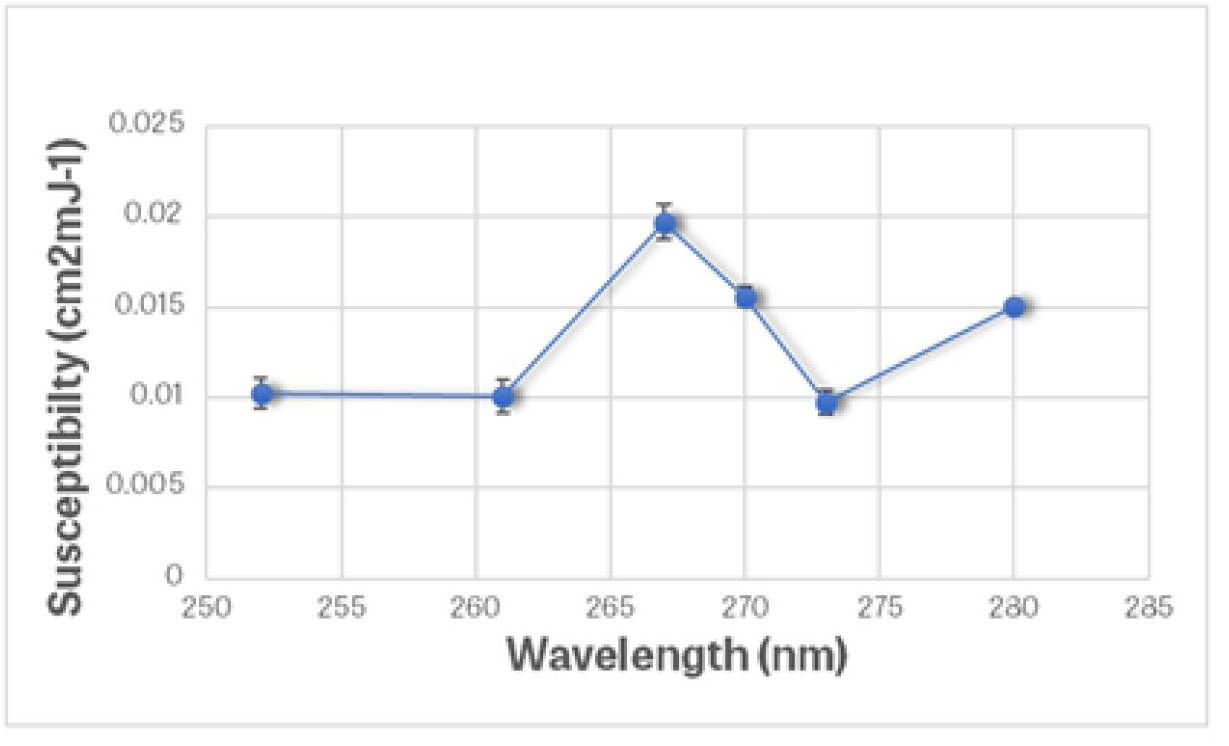
Susceptibility ([*k*] (cm^2^/mJ)) calculation to determine LRV obtained per mJcm^2^ against black mold (*Cladosporium halotolerans* ATCC 10391(NRRL 1671)). Error bars are given as the standard error of the linear regression.

Linear regression analysis at 95% for all UVC LEDs, 252 nm (R^2^ =0.9637, *p*=0.0030), 261 nm (R^2^ =0.9711, *p*=0.0021), 267 nm (R^2^ =0.9723, *p*=0.0020), 270 nm (R^2^ =0.9819, *p*=0.0010), 273 nm (R^2^ =0.9878, *p*=0.0006), and 280 nm (R^2^ =0.9914, *p*=0.0003) displayed significant association between peak wavelengths of the arrays and disinfection performances against *C. halotolerans* ATCC 10391.

Simulations were used towards designing and validating a strategy for preventing black mold growth on HVAC coils. The resulting minimum intensity at the outer limits of the coil are .12 mW/cm^2^ (120 µW/cm^2^) (Figure 5). This design would reach the target prevention dose in only four minutes of operation every twelve hours which provides significant annual energy savings over legacy mercury-containing lamp technology. In addition, there would not be any annual service fees or lamp replacement costs since the UVC LEDs would provide over 10 years of use.

**Figure 5:**
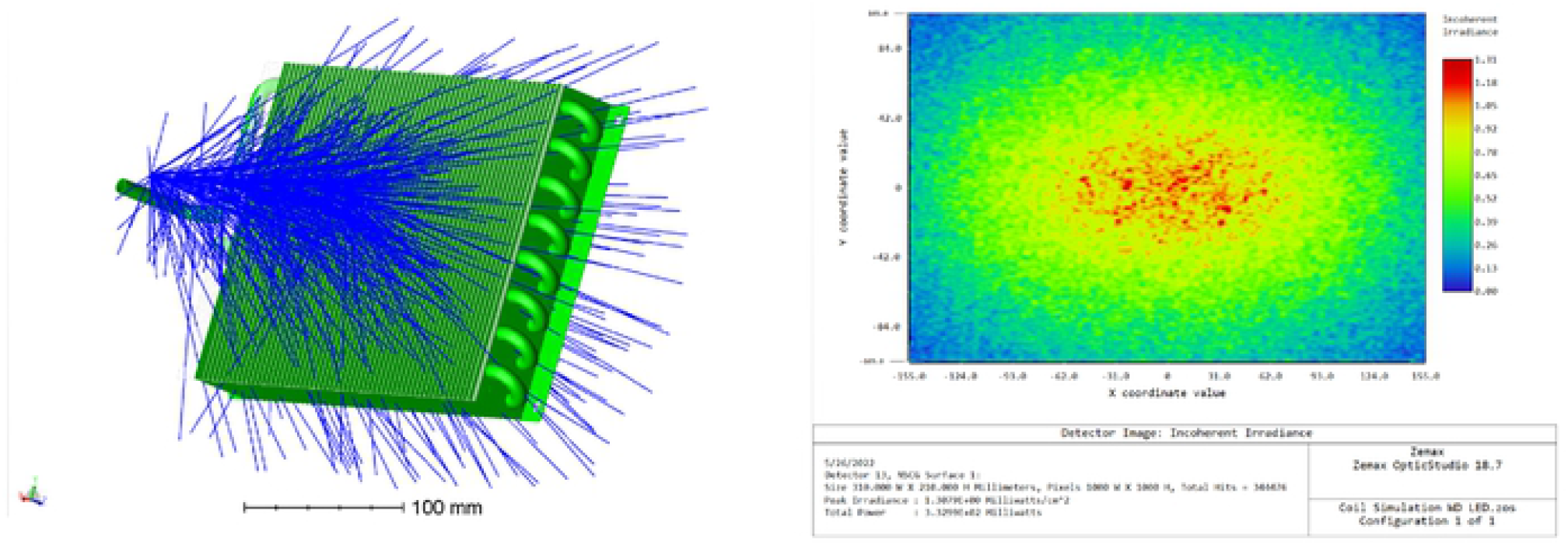
Simulation software can be used to determine the most efficient design based on dimensions. The use of UVC LEDs allows for the customization of variables such as power use, product lifetime, and costs in the product design phase.

Specifically, using periodic dosing at 267 nm peak wavelength, a low daily dose (28.8 mJ/cm^2^ every 12 hrs.) is enough to prevent mold growth and branching of hyphae on HVAC coils (Figure 6), whereas using static dosing, a higher dose (225 mJ/cm^2^) is needed to disinfect *C. halotolerans* to obtain 4.03 LRV (99.99% mold reduction). In general, inactivation of black mold requires more dose and thus best approach is to dose periodically to prevent mold from establishing in the first place [24]. Additionally, UVC disinfection is absorption-based and depends on the susceptibility of microbe genetic material to a particular UVC wavelength [25]. Incubation was done at 24°C and 97% RH. Genera mold confirmation using bright microscopy confirmed the mold type (Figure 2).

**Figure 6:**
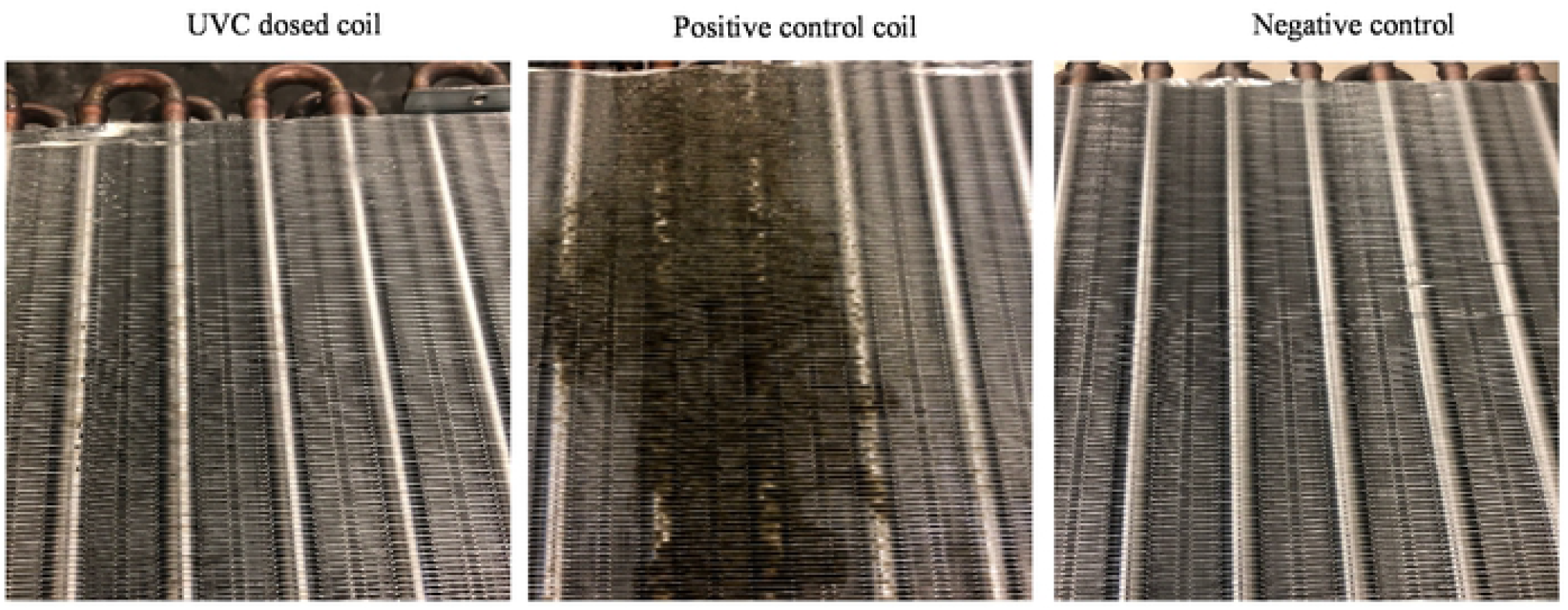
Coil mold disinfection using 9 LED array at 267 nm. No mold with periodic dosing for 4 min every 12 hrs. (28.8 mJ/cm^2^ every 12 hrs.)

## Discussion

The United Nations Environment Programme (UNEP) approved an international treaty to maintain public health and protect the environment from mercury pollution [26]. Luckily, new alternative UVC emission technologies such as LEDs now exist and are commercially available for use. The availability is in the form of different wavelengths, which influence performance, with peak wavelengths of around 265 nm offering better inactivation efficacies [27] due to higher germicidal efficacy [6]. One study that utilized an *E. coli* model organism found that 267 nm UVC LEDs have the highest inactivation efficiency compared to other studied wavelengths such as 275 and 310 nm [28]. Additionally, 267 nm, has been found to offer better performances against antibiotic-resistant *Bacillus* species [29], which are spore-forming bacteria just like black mold. The current study revealed that with 225 mJ/cm^2^ 4 LRV was obtained against *C. halotolerans* ATCC 10391 (Table 1). *C. halotolerans* is less susceptible compared to previously studied bacteria strains such as MRSA, *A. baumannii* and vancomycin-resistant *Enterococcus faecium* [30]. Like in the previous study, due to the inaccessibility of some LED packages, the 252 and 261 nm UVC LEDs were in packages with ball lenses, explaining their overall performance [31]. The relatively high germicidal efficacy of 267 nm is in alignment with previously observed trends in HVAC applications [6].

While investigating other simulations it was determined that as few as 4 LEDs per board could treat the side of a standard 14” coil found in most 3–5-ton residential HVAC units. LED numbers and operating conditions could be varied by manufacturers to provide lower-cost units with 2–4-year replacement cycles where recurring revenue is a consideration.

Experiments on HVAC coils to study mold growth prevention were done with relative humidity maintained at 97% (Figure 6). This offers conservative data if other molds were to be used since *Cladosporium halotolerans* survives humidity dynamics noticeably well in comparison to *Aspergillus niger* and *Penicillium rubens* [22]. Additionally, there is a reduced microbial susceptibility against UVC at high humidity [32]. However, the worst-case scenarios to test UVC LEDs should involve conditions such as high temperature, high relative humidity, and high salt concentrations, all of which encourage mold growth [33]. Because there is so little research on the susceptibility of fungi to UVC by wavelength, it’s difficult to compare results. However, these results do imply that *C. halotolerans* is slightly less susceptible to UVC than *A. fumigatus* as previously reported [34] which is consistent with other results [35], showing slightly more resistance of the *Cladosporium* genus. Because many studies do not report the fluence rate on the petri dish (but instead lamp power and time) it is difficult to know for sure, but this susceptibility is on the same order that was previously reported for *C. herbarum* [36].

Optimum duty cycle, period, and dose in the periodic dosing strategy were not explored because those optimums are expected to depend on environmental factors that will be difficult for designers to control. Susceptibility and growth rate are expected to depend on temperature, humidity, and surface conditions. These conditions are seldom constant over prolonged time periods in enclosed spaces [37]. As a result, varying these conditions to determine their effect was not considered for this study, and only went to high relative humidity (97%) to obtain conservative data. To maintain performance in enclosed spaces, it is expected integrators will implement a strategy to compensate for variation in conditions to ensure desired performance. The pulsed strategy using UVC LEDs has been found to be an attractive candidate for use in disinfection against bacteria [38], and it has now also been shown to prevent black mold growth and biofilms in HVAC systems.

## Conclusions

The study demonstrated that with a UVC dose of 225 mJ/cm2, 4.03 LRV (99.99 mold reduction) is obtained with a 267 nm array against *C. halotolerans* ATCC 10391. This study further confirms prior studies showing that commercially available UVC LEDs can be used as effective tools for disinfecting HVAC systems. Additionally, the study demonstrated that with a constant periodic dose of 28.8 mJ/cm^2^ every 12 hours prevention of mold growth on HVAC coils is possible. Since typical HVAC installation procedures include an initial cleaning of the coils, the pulsed dosing strategy is the most probable use case. Implementation of UVC LED devices might not only improve air quality, but due to prevention of mold growth in HVAC systems, could be central to energy cost savings by more than 25% due to periodic dosing, and could help people stay healthy in enclosed spaces.

## Abbreviations

ATCC: The American Type Culture Collection
CFU: Colony Forming Unit
HVAC: Heating, Ventilation, and Air Conditioning
LRV: Log Reduction Value
NRRL: Northern Regional Research Laboratory
PBS: Phosphate Buffered Solution
UVT: Ultraviolet Transmittance
UVC: Ultraviolet Subtype
C YM: Yeast Malt
RH: Relative Humidity

## Authors’ contributions

R.M.M. and R.V.R. designed the experiments. R.M.M. developed the methodology and coordinated the study. B.B. provided technical input during design of work and performed simulations. R.V.R. ensured provision and installation of systems for project. J.H.D. performed electrical measurements. R.M.M. and M.M.L. collected experimental data. R.M.M. and J.H.D. performed data analysis. R.M.M. wrote the original manuscript. J.H.D., and M.M.L. contributed to the writing of manuscript. All authors read and approved the final manuscript.

## Acknowledgements

Authors thank Dr. Kevin Kahn, Dr. Sanjay Kamtekar, Dr. Jianfeng Chen (Jeff), Patrick Aigeldinger and Thomas Horgan for their help with reviewing the manuscript. Authors are also grateful to Britt Hafner, Amy Miller, Chris Scully, and Ian Bonesteel for their technical help and constructive discussions. Authors finally acknowledge American Type Culture Collection, ATCC Genome Portal, available at https://genomes.atcc.org (Accessed: 20^th^ May October 2021).

## Data availability

All data generated or analyzed during this study are included in this published article. Spectral analysis raw data can be accessed in FigShare via https://doi.org/10.6084/m9.figshare.19742089.v1. Inactivation rate constant (*k*) analyses can be accessed in FigShare via https://doi.org/10.6084/m9.figshare.19742059.v1. Mold prevention time lapse videos can be accessed in FigShare via https://doi.org/10.6084/m9.figshare.19733176.v1.

